# Chromosome-Level Genome Assembly of Southern Africa Mozambique Tilapia (*Oreochromis mossambicus*) using PacBio HiFi and Omni-C sequencing

**DOI:** 10.1101/2025.11.27.690999

**Authors:** Thendo Stanley Tshilate, Lucky Tendani Nesengani, Sinebongo Mdyogolo, Rae Marvin Smith, Annelin Henriehetta Molotsi, Tracy Masebe, Ntanganedzeni Mapholi

## Abstract

The Mozambique tilapia (*Oreochromis mossambicus*) is a species of significant ecological and economic importance in Southern Africa. However, urbanization and water management challenges have led to the species being classified as threatened by the International Union for Conservation of Nature. Despite its widespread distribution and significance as a key food source, the genetic architecture of Southern African *O. mossambicus* remains inadequately characterized. This gap in knowledge hinders efforts to conserve and manage the species effectively. This study presents a high-quality chromosome-level genome assembly for a South African *O. mossambicus* assembled using both PacBio HiFi and Dovetail Omni-C sequencing. The 1.09 Gb assembly is highly contiguous, with 99.4% of sequences anchored into 22 chromosomes and a BUSCO completeness of 99.5%. We annotated 27,544 protein-coding genes. This genome provides an essential resource for preserving native biodiversity and enabling genomic selection for traits like disease resistance and growth performance within the South African aquaculture context.

## Background & Summary

Tilapia (*Oreochromis*) is the general name for cichlid fish, indigenous in the Middle East and Africa with over 70 existing species. The genus can thrive in an extensive range of environmental conditions which makes these one of the most farmed fish worldwide, mostly in subtropical and tropical regions^1–3^. Globally, tilapia sector is experiencing growth which highlighting its significant importance for food security and economies specially in developing countries ^4^. With South African tilapia aquaculture industry, the sector is still underdeveloped accounting for less than 1% of total global production^5,6^. The predominantly cultivated species are the indigenous Mozambique tilapia (*O. mossambicus*) and the non-indigenous Nile tilapia (*O. niloticus*) ^7–11^.

The *O. mossambicus* is ecologically and legally more suited for farming in South Africa, especially where environmental law restricts the use of indigenous tilapias. It can withstand the fluctuating climatic conditions, water shortage and has good tolerance of salinity and poor water quality ^7,8,12^. These traits makes it an outstanding aquaculture species with low-input cost with potential to farm extensive environment both in brackish and saline waters. Although they grow slower than *O. niloticus* species, their ecological hardness, high reproductive effectiveness and flexibility make *O. mossambicus* a suitable candidate for selective breeding programs ^13^. It also holds important local significance as a genetic resource of characters that may become increasingly valuable in climate-resilient aquaculture systems ^14^.

Previously, research studies on *O. mossambicus* have progressed in areas of population structure, hybridization, and genetic diversity^9,15–18^. Several studies have reported hybridization between native *O. mossambicus* and invasive species such as *O. niloticus* and *O. mortimeri* using mitochondrial DNA and microsatellite markers ^15–18^. These studies discovered large scale of genetic introgression which result in erosion of the indigenous gene pool which potentially endangers wild *O. mossambicus* populations. These reports played a significant part in alerting conservation agencies and aquaculture breeders to the genetic risks of hybridization and the need to preserve pure genetic stocks. Additionally, studies on genetic diversity further supported these trends where Simbine et al.^17^ reported that populations of *O. mossambicus* in southern Mozambique are declining due to reduced genetic variation which is a consequence of population bottlenecks and excess heterozygosity. Similarly, Mashaphu et al.^14^ observed the same pattern in South African wild populations reporting high genetic differentiation between groups and low diversity within populations. This decline in genetic variation was directly linked to human factors such as habitat fragmentation which is caused by urbanization and the construction of dams ^6,19,20^.

Recently, significant progress has been achieved in assembling high-resolution genomes for tilapia species for both aquaculture and conservation purposes. For *O. niloticus*, Conte et al.^21^ assembled a high-quality genome uncovering structural variations and duplicated sex determination loci. Building on this, Etherington et al. (2023) effectively reconstructed the X and Y haplotypes using chromosome-level assembly of Egyptian Abbassa strain, offering fresh perspectives on the genomic architecture connected to sex. With the aim of facilitating genomic selection, a high-quality chromosome-level genome of the *O. niloticus* GIFT strain was assembled and reported more than 11 Mb of introgressed *O. mossambicus* DNA also revealing genes related to immunity and growth through comparative analysis ^22^. The genome of *O. aureus* (blue tilapia) was also assembled and predicted around 23,000 genes. Comparative analyses with *O. niloticus* revealed conserved antimicrobial genes, highlighting its value in molecular breeding for disease resistance and in biomedical research ^23^. For *O. mossambicus*, a chromosome-level genome was assembled using sampled sourced at the Pearl River Fishery Research Institute in China and revealed novel features of its sex-determination pathways ^2^.

A lack of a high-quality reference genome tailored for Southern Africa O. *mossambicus* remains a key gap despite notable advancements in genome assembly research of this species. The distinctive genetic composition of southern African *O. mossambicus* is overlooked by most of the current assembly, which are based on Asian strains or hybrids. Given that South African *O. mossambicus* have unique genetic traits^14^ and that further hybridization poses a threat to additional genetic erosion ^7^, this is an urgent problem. Therefore, there is a need to develop a reference genome that is regionally suited to aid the preservation of native biodiversity. This will provide genetic insight which will assist in implementing selective breeding programmes and enhance conservation initiatives. The current project provide a high-quality genome of South African *O. mossambicus* generated using both Pacific Biosciences (PacBio) HiFi and Dovetail genomics Omni-C technologies.

## Methods

### Ethics statement

#### Ethics Statement

All procedures were approved by the University of South Africa Ethics Committee (Reference Number: AREC-100818-024). The study was conducted in accordance with relevant guidelines and regulations. The collection and use of the specimen were further authorized by the national Department of Agriculture, Land Reform and Rural Development (Reference Number: 12/11/1/1/23 (6508 AC)) and the Limpopo Department of Agriculture. The specimen was sourced as a deceased animal and no live animal experimentation was conducted.

### Sample collection, DNA extraction and quality Assessment

A deceased adult female *O. mossambicus* was collected at Madzivhandila College of Agriculture, Limpopo Province for sampling and transported using ice to Inqaba Biotechnology laboratory. The tissue was stored in a-80°C prior processing to preserve DNA integrity. High molecular weight (HMW) DNA was extracted from muscle using the Nanobind® tissue kit from Pacific Biosciences (PacBio), following the manufacturer’s protocol. The DNA integrity was evaluated using a Qubit 2.0 Fluorometer (Thermo Fisher Scientific), the fragment length and integrity was verified with the Femto Pulse System (Agilent Technologies).

### HiFi SMRTbell Library Preparation and Sequencing

The HIFI library was prepared using the SMRTbell Prep Kit 3.0 (PacBio) following the manufacturer’s protocol. High molecular weight (HMW) DNA was sheared first to a target size range (15–20 kb) using a Megaruptor3^24^ operations then after concentrated and purified using a 1× ratio of SMRTBell cleanup beads. The sheared DNA was repaired for damaged DNA using repair and A-tailing at 37□°C for 30 min and 65□°C for 5 min followed by adapter ligation at 20□°C for 30 min. Thereafter, a nuclease treatment step was performed at 37□°C for 15 min to remove un-ligated DNA and adapters. The library was then purified and size-selected with 3.1X v/v AMPure PB beads and fragments less than 10 kb were removed using BluePippin (Sage Science). The final SMRTbell library with average size between 15–20 kb was assessed and then sequenced using the PacBio Sequel II system with Sequel II chemistry 2.0 on 8M SMRT Cells with 24-hour movie collection time and PacBio (Sequel II® and Sequel IIe). The sequence generated 33.13 Gb of data with an average read length of 17,973 bp resulting in 33X coverage.

### Omni-C Library Preparation and Sequencing

Library was prepared from 5 mg of tilapia spleen tissue using the Dovetail Omni-C kit protocol with minor modifications. The tissue was first disrupted by grinding it to a fine powder using liquid nitrogen. The tissue was then transferred to a tube containing PBS and crosslinked with disuccinimidyl glutarate (DSG) for 10 minutes, followed by a 10-minute crosslinking with formaldehyde. Following the removal of the crosslinking agent, the sample was filtered through cell strainers measuring 200 μm and 50 μm to eliminate big tissue particles. For effective chromatin fragmentation and create long-range cis connections, the crosslinked chromatin was in situ digested using an optimal dosage of DNase I. The cells were lysed with sodium dodecyl sulfate (SDS) to liberate chromatin fragments. Two significant improvements were made to the library preparation in the proximity ligation stage: (1) the input lysate was reduced to reduce contaminants, and (2) the intra-aggregate bridge ligation step was extended from a 2-hour reaction to an overnight reaction to increase ligation efficiency. In short, digested chromatin fragments were ligated to biotinylated bridge adapters, end-repaired, and captured onto Chromatin Capture Beads. After that, proximity ligation was used to process the adapter-ligated fragments. After that, proteins were hydrolysed, crosslinks were reversed, and DNA was isolated. A sequencing library compatible with Illumina was then created from the DNA. Before PCR amplification, streptavidin beads were used to precipitate the biotinylated fragments. The final library was sequenced on an Illumina NovaSeq 6000 platform and generated two million 2 × 150 bp read pairs to evaluate mapping quality, true cis/trans interactions, and library complexity. For chromosome-level assembly, the Omni-C library was sequenced to approximately 100 million 2 × 150 bp read pairs per genome giga base (Gb) size producing produced 16.96 Gb of data.

### Genome Assembly

The HiFi and Omni-C sequencing data were put through several quality control and assembly procedures to generate a chromosome-level genome assembly for *O. mossambicus*. First, fastplong (v0.3.0) ^25^ was applied to raw reads from HiFi data to eliminate polyG tails, trim adapter sequences, and remove low-quality reads. Following this, the HiFi dataset consisted of 1.84 million reads with average length of 17,973 bp totalling to 33.13 Gb of sequence data. The Omni-C sequencing data underwent quality assessment using fastp (v0.24.0) ^26^, revealing high-quality paired-end reads (151 bp × 2), with 97.2% (113.08 million read pairs) passing filters. Genome structure was characterized through k-mer profiling using a k-mer size of 21. The Genomescope analysis estimated a genome size of 0.99 Gb and revealed a highly homozygous genome with a heterozygosity rate of 0.428% (homozygosity: 99.6%) (Figure 1).

**Figure 1.**
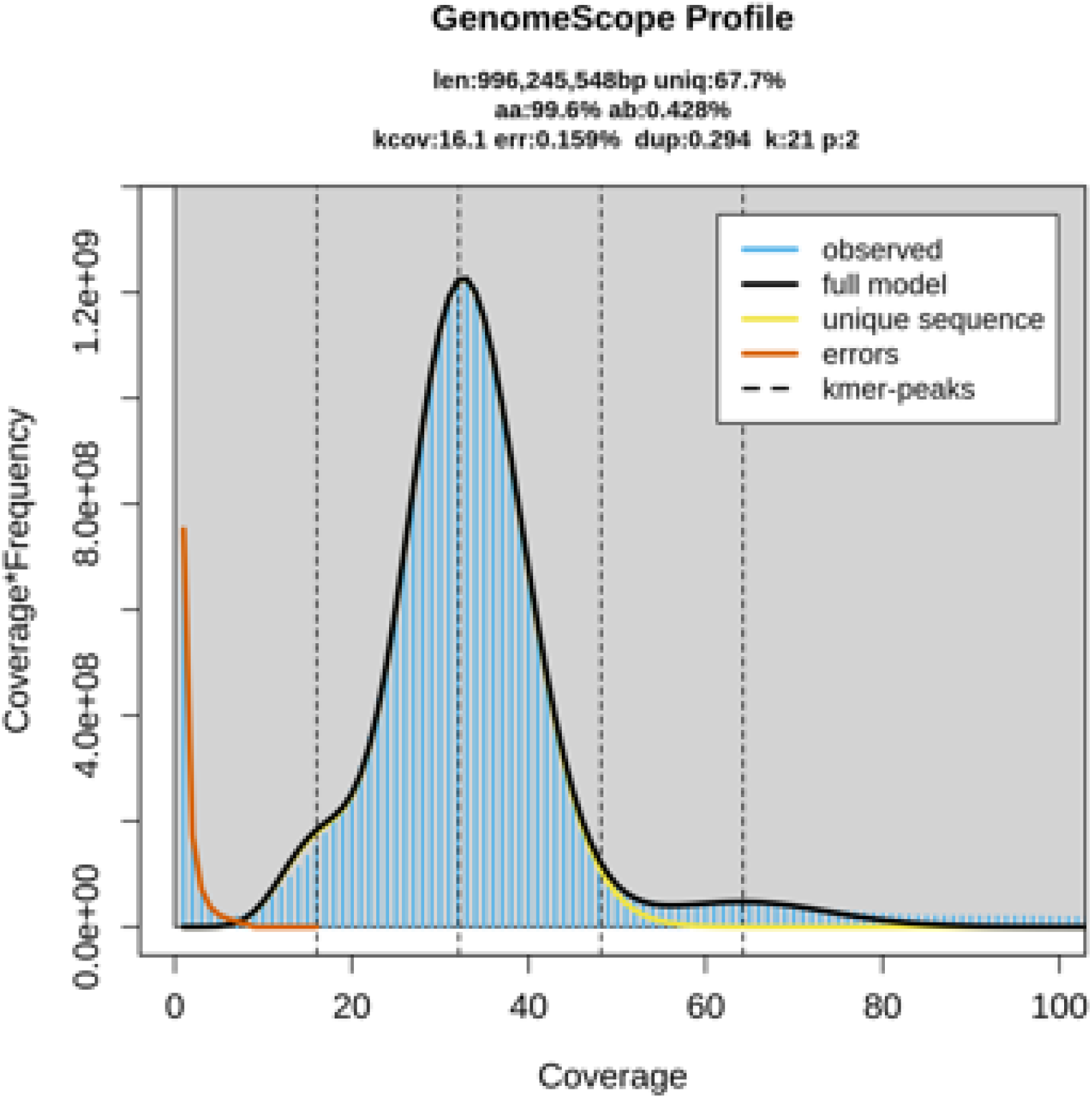
Genome size assessment of O. mossambicus by k-mer frequency estimated using Jellyfish and Genomescope

The genome was assembled with both PacBio HiFi reads and Dovetail Omni-C data using Hifiasm (v0.25.0)^27^ running in integrated Hi-C mode to produce a haplotype-resolved assembly. Omni-C workflow (https://dovetail-analysis.readthedocs.io/en/latest/index.html) from Dovetail Genomics, which combines several tools for contact map creation and library quality checking was used to process the Omni-C reads. Samtools was used to index the assembly and thereafter utilised to generate the “genome” file for further analysis. A high-quality, chromosome-scale genome assembly was generated through a multi-step process. First, Omni-C reads were aligned to the draft reference genome using BWA (v0.7.17)^28^, retaining only high-quality mapped reads. These reads were then processed with Pairtools (v1.0.3)^29^ to remove PCR duplicates and classify valid ligation events. The processed alignments were used to generate a Hi-C contact matrix with Juicer Tools (v1.6)^30^ which was subsequently scaffolded with YaHS (v1.1)^31^ to generate chromosome-level genome assembly. Assembly generated resulted in 1.09 Gb genome size consisting of 203 scaffolds with an N50 of 42.81 Mb. The assembly is highly contiguous, with 99.4% of the scaffolds correspond to 22 chromosomal blocks, representing the full diploid set of *O. mossambicus* chromosomes (Figure 2a, 2b; Table 1). The remaining 181 scaffolds are small, unmapped sequences. To assess completeness, the genic regions was analysed with BUSCO^32^ using the actinopterygii_odb10 database. The assembly captured 99.5% of the conserved orthologs (Figure 2c), a result comparable to previously published assemblies of related tilapia species (*O. niloticus* and *O. aureus*; Table 1). Further validation with k-mer analysis revealed a k-mer completeness of 95.5% and an indel quality value (QV) of 68.87 (Figure 2d), confirming the high base-level accuracy and contiguity of the assembled genome.

**Table 1.** Genome assembly statistics for *O. mossambicus* and other tilapia species.

|  |  |  |  |  |  |
| --- | --- | --- | --- | --- | --- |
|  | *O. mossambicus* | *O. mossambicus* | *O. niloticus* | *O. niloticus (GIFT)* | *O. aureus* |

|  |  |  |  |  |  |
| --- | --- | --- | --- | --- | --- |
| Sequencing Platform | PacBio, Omni-C | Illumina | Nanopore, Hi-C | Nanopore, Hi-C | Nanopore, Hi-C |
| Assembly level | Chromosomal | Scaffold | Chromosome | Scaffold | Chromosome |
| Genome size (Gb) | 1,09 | 1 | 1 | 1,1 | 1 |
| Chromosome Number | 22 | - | 22 | - | 22 |
| Scaffold Number | 203 | 51491 | 2459 | 23 | 303 |
| Scaffold N50 (Mb) | 42,81 | 1,5 | 30,8 | 41,9 | 40,7 |
| Scaffold L50 | 10 | 146 | 11 | 10 | 10 |
| Number of contigs | 264 | 85971 | 3009 | 406 | 1025 |
| Contig N50 | 35,6 | 0,0299 | 2,9 | 18,5 | 4,2 |
| Contig L50 | 13 | 8381 | 96 | 18 | 65 |
| GC percentage (%) | 40.9 | 40,5 | 40,5 | 41 | 40,5 |

**Figure 2.**
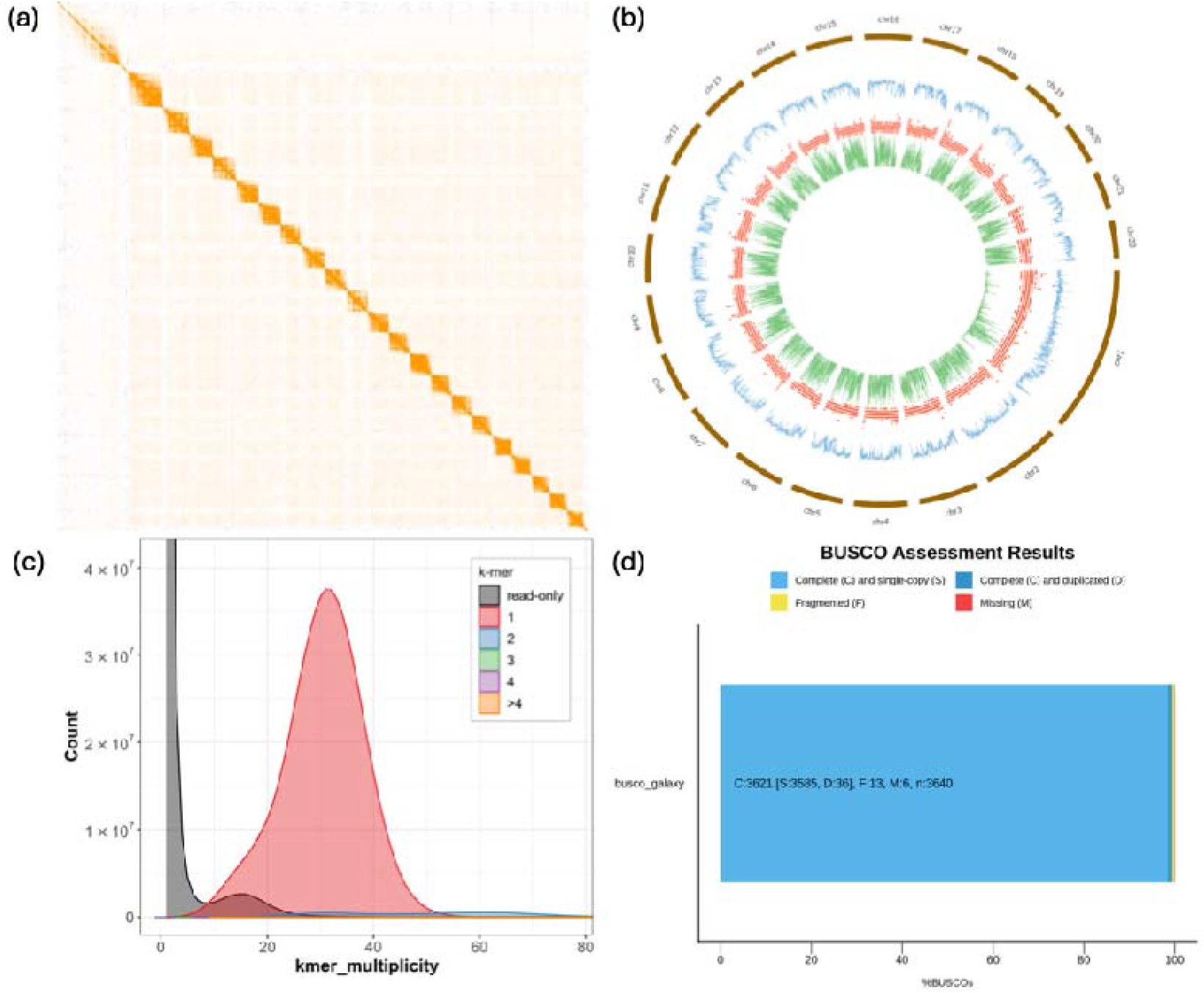
(a) Heatmap, pairwise interactions among 22 pseudochromosomes of *O. mossambicus* genome. colour depth represent the density of the Hi-C interactions (b) Genome coordinates and annotation of genome, blue colour represents the GC content, red represent repetitive elements and green represent the genes across the genome (c) Merqury assembly spectrum plots for evaluating k-mer completeness of *O. mossambicus* (d) BUSCO assessment results of *O. mossambicus* genome at the chromosome level.

### Genome annotation and functional characterisation

Repetitive elements were identified and soft-masked prior to gene annotation. Firstly, *de novo* repeat library was constructed using RepeatModeler v2.0.0, which integrates RepeatScout v1.0.6 ^33^, RECON v1.5.0 ^34^, and Tandem Repeats Finder (TRF) v4.09 ^35^. This library, along with known repeats from RepBase, was used as input for RepeatMasker v4.0.9 ^36^ to comprehensively annotate repetitive sequences. Identified repeats were subsequently annotated and classified using the “one_code_to_find_them_all” Perl script ^37^, providing a comprehensive characterization of transposable elements and other repetitive sequences in the genome. Repetitive sequences constitute 43.78% (483 Mb) of the *O. mossambicus* genome. Long interspersed nuclear elements (LINEs) were the most abundant class of known repeats, accounting for 8.71% of the genome. A full breakdown of repetitive elements is provided in Table 2.

**Table 2.** Summary of repeat elements identified in the genome of the *O. mossambicus*.

| Repeat element | Number of elements | Length (bp) | Percentage(%) |
| --- | --- | --- | --- |
| SINEs | 364227 | 143588361 | 13,01 |
| LINEs | 237619 | 96139142 | 8,71 |
| LTR elements | 76968 | 33837694 | 3,06 |
| DNA element | 541264 | 145128603 | 13,15 |
| Small RNA | 40000 | 12368229 | 1,12 |
| Simple repeat | 362510 | 14823593 | 1,34 |
| Low complexity | 43190 | 2059482 | 0.19 |
| Unclassified | 694471 | 166274202 | 15,6 |

The masked genome was annotated using *ab initio* gene prediction tool TIBERius v1.1.4 ^38^. This tool integrates convolutional and long short-term memory (LSTM) layers with a hidden Markov model (HMM) to predict genes with high accuracy directly from genomic sequence. A key strength of TIBERius lies in its ability to predict gene structures directly from genomic sequences with high accuracy, comparable to annotation pipelines that rely on extrinsic evidence such as RNA-seq or protein homology data. A total of 27,544 protein-coding genes were identified with each with a single associated transcript, indicating a one-to-one correspondence between genes and their primary isoforms (Table 3). The average gene length was 10,298 bp ranging from 201 bp to 588,982 bp.

**Table 3.** Gene structure and gnome annotation results for *O. mossambicus*.

|  |  |
| --- | --- |
| Number of genes | 27544 |
| Mean gene length (bp) | 10298.4 |
| Number of exons per gene | 8.00218 |
| Mean exon length (bp) | 176.466 |
| Percentage of genes with introns | 87.195 |
| Mean intron length (bp) | 1269.07 |
| Total intron length (Mb) | 244.764 |
| Average transcript length (bp) | 10298.4 |

For functional annotation, proteins predicted were analysed using eggNOG-mapper v2.1.12, which assigns annotation information of ortholog from the eggNOG database ^39^. This assigned putative functions to 21,229 genes (77%) based on homology to orthologous groups and matches to multiple protein databases, including Gene Ontology (GO), Kyoto Encyclopaedia of Genes and Genomes (KEGG), Pfam, CAZy, BRITE, and BiGG. (Figure 3 and Supplementary Table 1).

**Figure 3.**
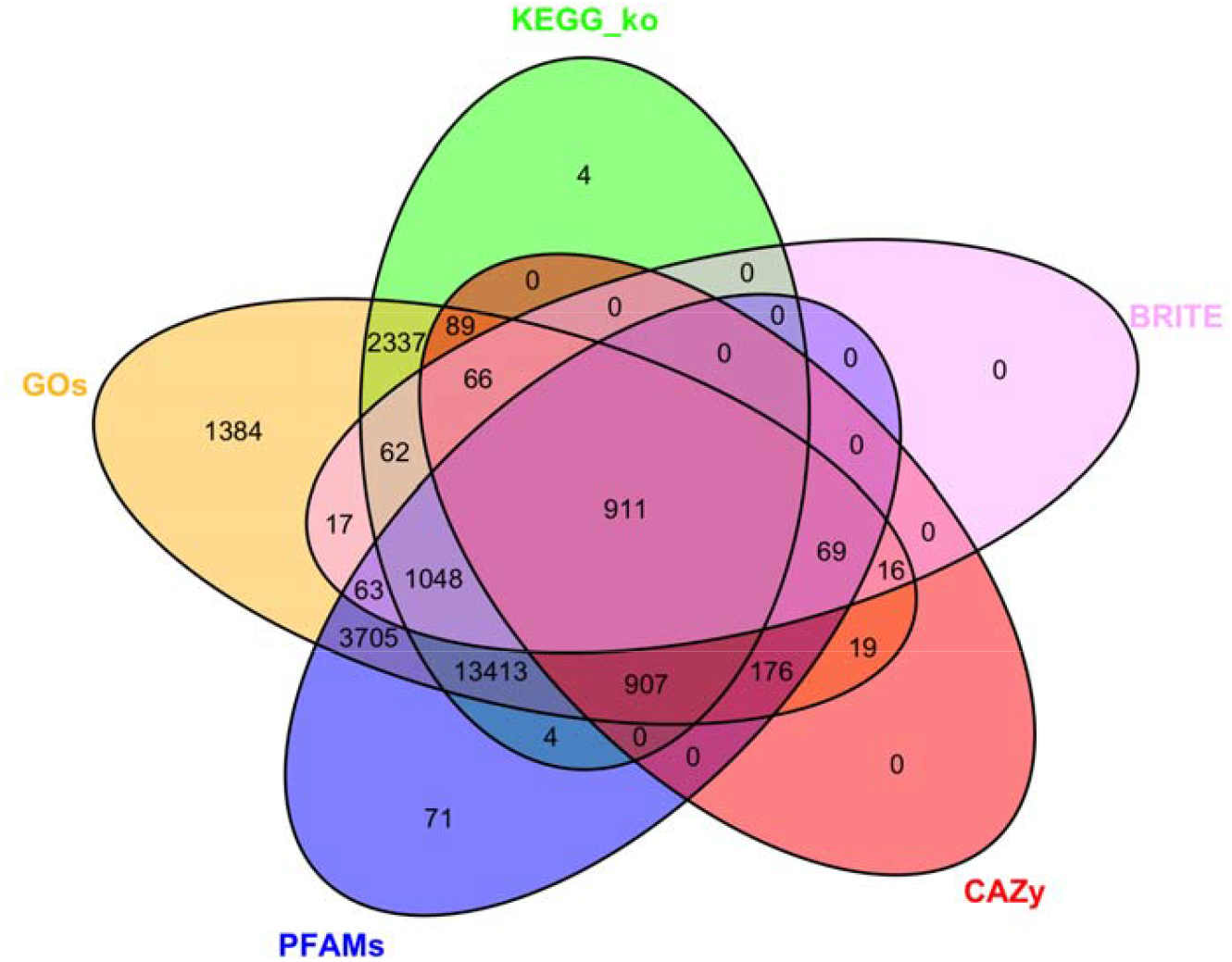
Venn diagram of function annotations from various database displaying the overlap and uniqueness of functional gene annotation derived from five databases.

## Data availability

The sequencing data for this study have been deposited in the NCBI under BioProject PRJNA1227266, with Sequence Read Archive (SRA) accessions SRR35773053 and SRR35773052. The genome assembly is available at NCBI under the submission ID SUB15704679 and will be accessible through the Genome database upon completion of NCBI processing. For reviewers, the genome assembly and annotation files are accessible via Figshare: https://doi.org/10.6084/m9.figshare.30366010

## Conflict of interest

The authors declare no conflict of interest.

### Technical Validation

High-quality genomic DNA was assessed before SMRTbell library preparation to ensure sufficient concentration and integrity for long-read sequencing. DNA concentration was measured using Qubit Fluorometer (Thermo Fisher Scientific) with the Qubit dsDNA BR and HS Assay Kits following the manufacturer’s protocol. A Qubit working solution was prepared by mixing 1 µL of fluorescent dye with 199 µL of buffer. For each measurement, a 1 µL of DNA sample was added to 199 µL of working solution then thoroughly mixed and incubated for 2 minutes at room temperature before measurement. DNA concentration (ng/µL) was automatically calculated by the Qubit software. The integrity size of the DNA was evaluated with Agilent Femto Pulse system with the Genomic DNA 50 kb kit (HS 50 kb). A 2 µL of genomic DNA was assed to 22 µL of loading dye following manufacturer’s protocol. This analysis provided an electropherogram and DNA size distribution to confirm high molecular weight DNA suitable for SMRTbell library preparation and also specify the final library size to be loaded for sequencing.

## Code availability

Software and pipelines used in this study were executed following the official manuals and published protocols of the respective bioinformatics tools. The specific software versions are provided in the Methods section.

## Acknowledgements

We acknowledge the University of South Africa for funding and support. This study forms part of the African BioGenome Project (AfricaBP): The Nelson Mandela Genomes Initiative for Conservation of Nature. We sincerely thank all the team at our research group for their guidance, project management and provision of resources. Finally we acknowledge Madzivhandila College of Agriculture for supplying the biological samples essential for this research.

## References

1. Agresti, J. J. et al. Breeding new strains of tilapia: Development of an artificial center of origin and linkage map based on AFLP and microsatellite loci. Aquaculture 185, 43–56 (2000).

2. Tao, W. et al. A Chromosome-Level Genome Assembly of Mozambique Tilapia (Oreochromis mossambicus) Reveals the Structure of Sex Determining Regions. Front Genet 12, 796211 (2021).

3. Lee, B. Y. et al. A Second-Generation Genetic Linkage Map of Tilapia (Oreochromis spp.). Genetics 170, 237 (2005).

4. FAO STATISTICS. https://www.fao.org/fishery/static/Yearbook/YB2019_USBcard/index.htm.

5. Mbokane, E. M. & Moyo, N. A. G. Use of medicinal plants as feed additives in the diets of Mozambique tilapia (Oreochromis mossambicus) and the African Sharptooth catfish (Clarias gariepinus) in Southern Africa. Front Vet Sci 9, 1072369 (2022).

6. Moyo, N. A. G. & Rapatsa, M. M. A review of the factors affecting tilapia aquaculture production in Southern Africa. Aquaculture 535, (2021).

7. D’Amato, M. E., Esterhuyse, M. M., Van Der Waal, B. C. W., Brink, D. & Volckaert, F. A. M. Hybridization and phylogeography of the Mozambique tilapia Oreochromis mossambicus in southern Africa evidenced by mitochondrial and microsatellite DNA genotyping. Conservation Genetics 8, 475–488 (2007).

8. Mutshekwa, T. et al. Behavioural Responses and Mortality of Mozambique Tilapia Oreochromis mossambicus to Three Commonly Used Macadamia Plantation Pesticides. Water (Switzerland) 14, 1257 (2022).

9. Mashaphu, M. F., O’Brien, G. C., Downs, C. T. & Willows-Munro, S. Genetic assessment of farmed Oreochromis mossambicus populations in South Africa. PeerJ 13, e18877 (2025).

10. Ansah, Y. B., Frimpong, E. A. & Hallerman, E. M. Genetically-improved tilapia strains in Africa: Potential benefits and negative impacts. Sustainability (Switzerland) 6, 3697–3721 (2014).

11. Wu, Y. rui et al. Effect of Sophora flavescens on non-specific immune response of tilapia (GIFT Oreochromis niloticus) and disease resistance against Streptococcus agalactiae. Fish Shellfish Immunol 34, 220–227 (2013).

12. Baroiller, J. F., D’Cotta, H., Bezault, E., Wessels, S. & Hoerstgen-Schwark, G. Tilapia sex determination: Where temperature and genetics meet. Comparative Biochemistry and Physiology - A Molecular and Integrative Physiology 153, 30–38 (2009).

13. Eknath, A. E. et al. Genetic improvement of farmed tilapias: the growth performance of eight strains of Oreochromis niloticus tested in different farm environments. Aquaculture 111, 171–188 (1993).

14. Mashaphu, M. F., Downs, C. T., Burnett, M., O’Brien, G. & Willows-Munro, S. Genetic diversity and population dynamics of wild Mozambique tilapia (Oreochromis mossambicus) in South Africa. Glob Ecol Conserv 54, e03043 (2024).

15. D’Amato, M. E., Esterhuyse, M. M., Van Der Waal, B. C. W., Brink, D. & Volckaert, F. A. M. Hybridization and phylogeography of the Mozambique tilapia Oreochromis mossambicus in southern Africa evidenced by mitochondrial and microsatellite DNA genotyping. Conservation Genetics 8, 475–488 (2007).

16. Sarker, B. S., Ali, A., Rahman, S. S., Alam, M. S. & Islam, M. S. Monogamous hybridization of Nile tilapia (Oreochromis niloticus) with Mozambique tilapia (O. mossambicus) results in unprecedented all-female F1 hybrid. Aquac Fish https://doi.org/10.1016/J.AAF.2024.04.007 (2024) xdoi:10.1016/J.AAF.2024.04.007.

17. Simbine, L., Viana da Silva, J. & Hilsdorf, A. W. S. The genetic diversity of wild Oreochromis mossambicus populations from the Mozambique southern watersheds as evaluated by microsatellites. Journal of Applied Ichthyology 30, 272–280 (2014).

18. Hong Xia, J. et al. Signatures of selection in tilapia revealed by whole genome resequencing. Sci Rep 5, 1–10 (2015).

19. Adom, R. K. & Simatele, M. D. The role of stakeholder engagement in sustainable water resource management in South Africa. Nat Resour Forum 46, 410–427 (2022).

20. Mirante, D. et al. Fine-tuning coexistence: Wildlife’s short-term responses to dynamic human disturbance patterns. Glob Ecol Conserv 54, e03053 (2024).

21. Conte, M. A., Gammerdinger, W. J., Bartie, K. L., Penman, D. J. & Kocher, T. D. A high quality assembly of the Nile Tilapia (Oreochromis niloticus) genome reveals the structure of two sex determination regions. BMC Genomics 18, 1–19 (2017).

22. Etherington, G. J. et al. Chromosome-level genome sequence of the Genetically Improved Farmed Tilapia (GIFT, Oreochromis niloticus) highlights regions of introgression with O. mossambicus. BMC Genomics 23, 1–16 (2022).

23. Bian, C. et al. Whole Genome Sequencing of the Blue Tilapia (Oreochromis aureus) Provides a Valuable Genetic Resource for Biomedical Research on Tilapias. Mar Drugs 17, 386 (2019).

24. Bates, A., Clayton-Lucey, I. & Howard, C. Sanger Tree of Life HMW DNA Fragmentation: Diagenode Megaruptor®3 for LI PacBio v1. https://doi.org/10.17504/PROTOCOLS.IO.81WGBXZQ3LPK/V1 (2023) xdoi:10.17504/PROTOCOLS.IO.81WGBXZQ3LPK/V1.

25. Chen, S. Ultrafast one-pass FASTQ data preprocessing, quality control, and deduplication using fastp. iMeta 2, e107 (2023).

26. Chen, S., Zhou, Y., Chen, Y. & Gu, J. Fastp: An ultra-fast all-in-one FASTQ preprocessor. Bioinformatics 34, i884–i890 (2018).

27. Cheng, H., Concepcion, G. T., Feng, X., Zhang, H. & Li, H. Haplotype-resolved de novo assembly using phased assembly graphs with hifiasm. Nat Methods 18, 170–175 (2021).

28. Li, H. Aligning sequence reads, clone sequences and assembly contigs with BWA-MEM. 00, 1–3 (2013).

29. Open2C et al. Pairtools: from sequencing data to chromosome contacts. bioRxiv 2023.02.13.528389 (2023) doi:10.1101/2023.02.13.528389.

30. Durand, N. C. et al. Juicer Provides a One-Click System for Analyzing Loop-Resolution Hi-C Experiments. Cell Syst 3, 95–98 (2016).

31. Zhou, C., McCarthy, S. A. & Durbin, R. YaHS: yet another Hi-C scaffolding tool. Bioinformatics 39, (2023).

32. Tegenfeldt, F. et al. OrthoDB and BUSCO update: annotation of orthologs with wider sampling of genomes. Nucleic Acids Res 53, D516–D522 (2025).

33. Price, A. L., Jones, N. C. & Pevzner, P. A. De novo identification of repeat families in large genomes. BIOINFORMATICS 21, 351–358 (2005).

34. Bao, Z. & Eddy, S. R. Automated de novo identification of repeat sequence families in sequenced genomes. Genome Res 12, 1269–1276 (2002).

35. Benson, G. Tandem Repeats Finder: A Program to Analyze DNA Sequences. Nucleic Acids Research vol. 27 https://academic.oup.com/nar/article/27/2/573/1061099 (1999).

36. Tarailo-Graovac, M. & Chen, N. Using RepeatMasker to identify repetitive elements in genomic sequences. Curr Protoc Bioinformatics Chapter 4, (2009).

37. Bailly-Bechet, M., Haudry, A. & Lerat, E. ‘One code to find them all’: A perl tool to conveniently parse RepeatMasker output files. Mob DNA 5, 1–15 (2014).

38. Gabriel, L., Becker, F., Hoff, K. J. & Stanke, M. Tiberius: End-to-End Deep Learning with an HMM for Gene Prediction. bioRxiv 2024.07.21.604459 (2024) doi:10.1101/2024.07.21.604459.

39. Cantalapiedra, C. P., Hern[andez-Plaza, A., Letunic, I., Bork, P. & Huerta-Cepas, J. eggNOG-mapper v2: Functional Annotation, Orthology Assignments, and Domain Prediction at the Metagenomic Scale. Mol Biol Evol 38, 5825–5829 (2021).

